# Integrating past, present and future wheat research with Pretzel

**DOI:** 10.1101/517953

**Authors:** Gabriel Keeble-Gagnère, Don Isdale, Rados-ław Suchecki, Alex Kruger, Kieran Lomas, David Carroll, Sean Li, Alex Whan, Matthew Hayden, Josquin Tibbits

## Abstract

**Motivation:** Major advances have been made in the assembly of complex genomes such as wheat. Remaining challenges include linking these new resources to legacy research, as well as lowering the bar to entry for members of the research community who may lack specific bioinformatics skills.

**Results:** Pretzel aims to solve these problems by providing an interactive, online environment for data visualisation and analysis which, when loaded with appropriately curated data, can enable researchers with no bioinformatics training to exploit the latest genomic resources. We demonstrate that Pretzel can be used to answer common questions asked by researchers as well as for advanced curation of pseudomolecule structure.

**Availability:** Pretzel is implemented in JavaScript and is freely available under a GPLv3 license online at: https://github.com/plantinformatics/pretzel.

## 1 Introduction

The release of the RefSeq v1.0 genome assembly of bread wheat cultivar Chinese Spring (International Wheat Genome Sequencing Consortium, 2018) provided the first detailed map of the 16Gb genome bringing wheat genomic resources closer to the standard enjoyed in rice and maize. Already, international efforts are in the process of assembling more than ten elite wheat varieties to a similar standard (10+ Wheat Genomes Project, http://www.10wheatgenomes.com/). Within five years we can expect hundreds of high quality wheat genomes to be available. The challenge that remains is to connect these new resources to the wealth of research carried out in the past few decades, and make this information accessible to researchers and breeders who may lack the required bioinformatics skill-set. Existing tools such as cMap (Fang *et al.*, 2003) or genome browsers (Stein, 2013; Buels *et al.*, 2016) allow the visualisation of genetic maps and genome assemblies, respectively, but are separate tools. Hence one cannot view a genetic map alongside a physical genome sequence, for example. Wheat-focused databases such as GrainGenes (https://wheat.pw.usda.gov/GG3/) and T3 (Blake *et al.*, 2016) provide a wealth of data but are primarily repositories rather than environments for interactive visualisation. Pretzel aims to provide a framework that combines curated data stored in a back-end with a modern front-end graphical user interface for visualisation and analysis via a web browser.

## 2 Methods

Pretzel is built on several JavaScript frameworks: the front-end combines Ember.js with D3.js for real-time visualisation; the back-end uses Loopback.js connecting to a MongoDB database. The user interacts with Pretzel via a web page served by the front-end. The data model has been designed to be as general as possible while still enabling the richness of data to be captured. The highest level data structure is a *dataset*, which contains one or more *blocks*. For example, a dataset may describe a genetic map, where the blocks are individual linkage groups. Within each block are a set of *features* which are defined as intervals (possibly of zero length, which would define a single position) within the block. For example, a feature could be a molecular marker in a linkage group, or the position of a gene in a physical chromosome. Using only these three concepts, a large number of concepts can be represented: a quantitative trait locus (QTL) is simply an interval *feature* within a linkage group *block*; whereas pseudomolecule structure can be described by a set of intervals denoting scaffold start/end positions *features* within a chromosome *block*. As there are often multiple alternative names for molecular markers we created a fourth concept *alias*. *Aliases* enables features with different names to be considered as the same and therefore linked when drawing alignments. At present this general concept allows the association of syntenic genes between genomes and the direct comparison of genetic maps based on different marker systems/types through alias by reference position.

Data upload is simple and can be achieved through the web interface in CSV or JSON format (the native data structure in JavaScript); or via command line in JSON format directly to the back-end. Data sets can be uploaded in whole or in part, or as they are generated, providing flexibility and allowing users to add data in real time. Upon upload, data is visible to that user only, unless they explicitly make it available to other users. In addition, we have created a pipeline, https://github.com/plantinformatics/pretzel-input-generator), to generate Pretzel-ready data from standard genome assembly formats in an automated way. This pipeline has been used to make available a set of publicly available genomes so users can quickly set up a Pretzel server populated with real data.

A common goal in wheat genetic studies is to fine map a QTL to discover the underlying gene. This generally requires the translation of the QTL region from genetic map space to physical genome space. Once this is achieved one of the most common questions asked in wheat research is to ask what genes underlie a QTL interval defined in a genetic mapping experiment. With Pretzel, users with no bioinformatics training can quickly project their genetic map interval onto physical sequence and extract the genes in the target region for downstream analysis. An additional challenge when fine mapping an interval is potential mis-assembly in the region of interest. Figure 1 shows a genetic map aligned to a physical chromosome, as well as two syntenic chromosomes (chromosomes 1B and 1D) aligned via their syntenic genes (shown in red). By comparing the co-linearity of genes between sub-genomes, potential assembly problems can be flagged by noting when abrupt changes in co-linearity occur at scaffold break points. In this way, advanced curation of pseudomolecule structure can be achieved on any number of genomes.

**Figure 1:**
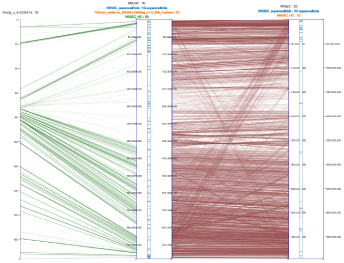
Pretzel alignment showing a 90k genetic map on the left against the IWGSC RefSeq v1.0 chromosome 1B pseudomolecule in the centre, with chromosome 1D on the right. Green lines between features indicate direct, e.g. marker-based links. Red lines indicate links via alias which can be defined e.g. based on sequence similarity between genes. The two pseudomolecules have their scaffold arrangement (AGP structure) displayed in blue, alongside their axes.

The technology stack Pretzel is built on is lightweight and a user with the necessary dependencies can clone, build and run the Pretzel code within minutes with a few simple commands. If run on an internet-facing machine, external users can then connect to the server. To make the installation process even easier, we provide Docker container images on Docker Hub (https://hub.docker.com/r/plantinformaticscollaboration/pretzel/). An instance of Pretzel populated with Australia-specific data is available to Grains Research and Development Corporation-funded researchers at http://plantinformatics.io/.

## 3 Conclusion

We have developed a web-based visualisation and analysis tool that solves some of the challenges present today in wheat research. Its extensibility and flexibility provides a strong foundation for future development of new features. The scalability of its technology stack ensures that Pretzel will accommodate the volumes of data that will become available in the next few years.

## References

Blake, V. C., Birkett, C., Matthews, D. E., Hane, D. L., Bradbury, P., and Jannink, J.-L. (2016). The triticeae toolbox: Combining phenotype and genotype data to advance small-grains breeding. 9. 2.

Buels, R., Yao, E., Diesh, C. M., Hayes, R. D., Munoz-Torres, M., Helt, G., Goodstein, D. M., Elsik, C. G., Lewis, S. E., Stein, L., and Holmes, I. H. (2016). Jbrowse: a dynamic web platform for genome visualization and analysis. Genome Biology, 17(1), 66.

Fang, Z., Polacco, M., Chen, S., Schroeder, S., Hancock, D., Sanchez, H., and Coe, E. (2003). cmap: the comparative genetic map viewer. Bioinformatics, 19(3), 416-417.

International Wheat Genome Sequencing Consortium (2018). Shifting the limits in wheat research and breeding using a fully annotated reference genome. Science, 361(6403).

Stein, L. D. (2013). Using gbrowse 2.0 to visualize and share next-generation sequence data. Briefings in Bioinformatics, 14(2), 162-171.

